# Searching for Simple Rules in *Pseudomonas aeruginosa* Biofilm Formation

**DOI:** 10.1101/705541

**Authors:** William Deveaux, Kumar Selvarajoo

## Abstract

Living cells display complex and non-linear behaviors, especially when posed to environmental threats. Here, to understand the self-organizing cooperative behavior of a microorganism *Pseudomonas aeruginosa*, we developed a discrete spatiotemporal cellular automata model based on simple physical rules, similar to Conway’s game of life. The time evolution model simulations were experimentally verified for *P. aeroginosa* biofilm for both control and antibiotic azithromycin (AZM) treated condition. Our model suggests that AZM regulates the single cell motility, thereby resulting in delayed, but not abolished, biofilm formation. In addition, the model highlights the importance of reproduction by cell to cell interaction is key for biofilm formation. Overall, this work highlights another example where biological evolutionary complexity may be interpreted using rules taken from theoretical disciplines.

## 1. Introduction

Under external stress, microorganisms, such as bacteria and fungi, are able to produce cooperative response by forming biofilm. Biofilm are aggregates of cells, often produced through quorum sensing mechanisms or due to the secretion of extracellular polymeric substance [1]. They often pose problems to food and water safety. On the other hand, biofilm caused by pathogenic agents in human often show drug resistance [2]. Thus, each year, billions of dollars are at risk due to biofilm-mediated damage [3]. Despite the immense research over decades using low to high throughput experimental methodologies, the progress in understanding the regulatory mechanisms or controlling the progression of biofilm is highly limited. Therefore, better knowledge in biofilm growth and evolution is necessary, and scientists could explore interdisciplinary strengths to fill the missing gaps.

The pathogenic Gram-negative bacteria *P. aeruginosa*, is well-known for its intrinsic and acquired antibiotic resistance. It facilitates chronic infections in human by its rapid ability to form biofilms [4]. Most previous works have mainly studied *P. aeruginosa* antibiotic resistance in their planktonic or single cell cultures. However, *P. aeruginosa* infection quickly spreads and form biofilm in certain diseases such as in the cystic fibrosis of the lung [5]. Thus, when infected in human, *P. aeruginosa* can cause death, especially for patients with cystic fibrosis, as they form biofilm that are resistant to current multi-drug antibiotic regimens [6]. Hence, the species and its biofilm are of considerable importance to medical care and patients’ well-being.

In this paper, we studied the self-organization of planktonic single cell stage to cooperative biofilm stage of *P. aeruginosa*. For understanding self-organization, a large number of theoretical and computational works uses continuous differential equation approaches [7,8], where the models require detailed mechanistic parameter values that are difficult to obtain from living systems. To overcome this limitation, here we utilized a discrete spatiotemporal computational methodology or cellular automata (CA), to predict the growth evolution mechanisms of *P. aeruginosa* [9]. CA models adopt simple physical rules or differential equations that matches experimental observation. By doing so, one can estimate the governing process or rules underlying the proliferation or cooperative behaviors.

Although there have been several efforts to model biofilm growth using CA in general [10,11], here we focused on simple rules to specifically test 2-D spatio-temporal dynamics of *P. aeruginosa* biofilm in untreated (control) and antibiotic treated conditions.

## 2. Data

For experimental observation, we obtained time-series growth profiles of *P. aeruginosa* performed by Gillis and Iglewski [6], see Fig. 1. Here, data for 2 conditions are shown; A) *without* and B) *with* 2 μg/ml of azithromycin (or AZM, a macrolide antibiotic). We observe that untreated *P. aeruginosa* rapidly forms clustered biofilm within 72h. The treated cells, although display biofilm emergence at 72h, shows a much slower formation time.

**Fig. 1.**
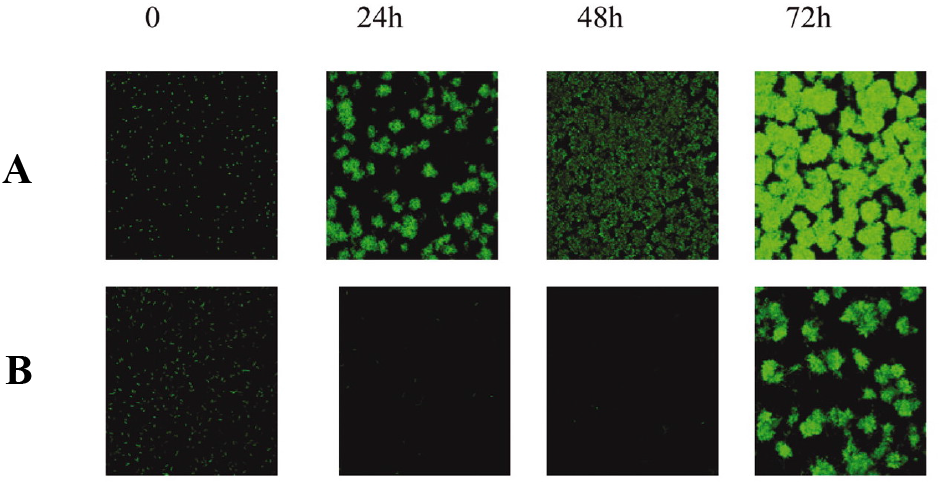
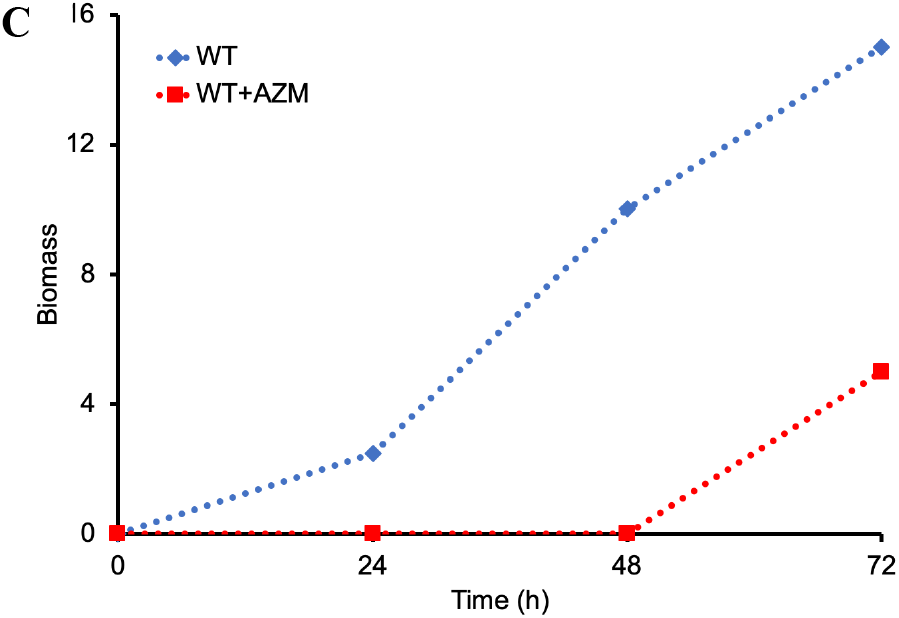
Time series confocal microscopy of *P. aeruginosa* PAO1 colony, (A) without and (B) with antibiotic (azithromycin or AZM) treatment. (C) Total biomass (μm^2^/μm^3^) in time. Note: biomass will be used as a proxy for cell numbers for simulations. Figures modified from [6], and permission to reproduce obtained.

## 3. Methods

Previously, to study cancer cell proliferation in control and drug treated conditions, we developed a discrete spatiotemporal CA model based on simple rules modified from Conway’s game of life [12,13]. The model’s rules were iteratively guessed and modified until the simulations matched experimental observations [13]. The resultant rules were used to infer the proliferation properties of the control and treated cancer cells. Here, we extended the model to predict the biofilm formation of *P. aeruginosa*.

### 3.1. Spatial Temporal Cellular Automata

#### 3.1.1 Cellular Automata Model

A 3-D CA model was developed in Matlab code consisting of 400 × 400 × 4 cubic (640,000) cells, with each cell having maximum 17 neighbors for the top and bottom planes, while 26 neighbors for other planes. The 8 corners, however, have a maximum of 7 neighbors, and 11 neighbors on the edges. We choose the empty initial cells large enough to avoid reaching the edges/corners within the simulated time steps. Note that our z-axis is small as the cells were originally cultured in 2-D flow cells plates, where cells often limited to a few layers on the vertical axis.

At time = 0h, for each condition, the cells were populated with live cells in random orientation that filled the spaces similar to Figure 1. We found 5,000 cells (Fig. 2A, leftmost panel) were distributed in a way that was similar to actual cell distribution in Figure 1. The CA rules (see subsection 3.1.2 below) were applied from time step 1b onwards.

**Fig. 2.**
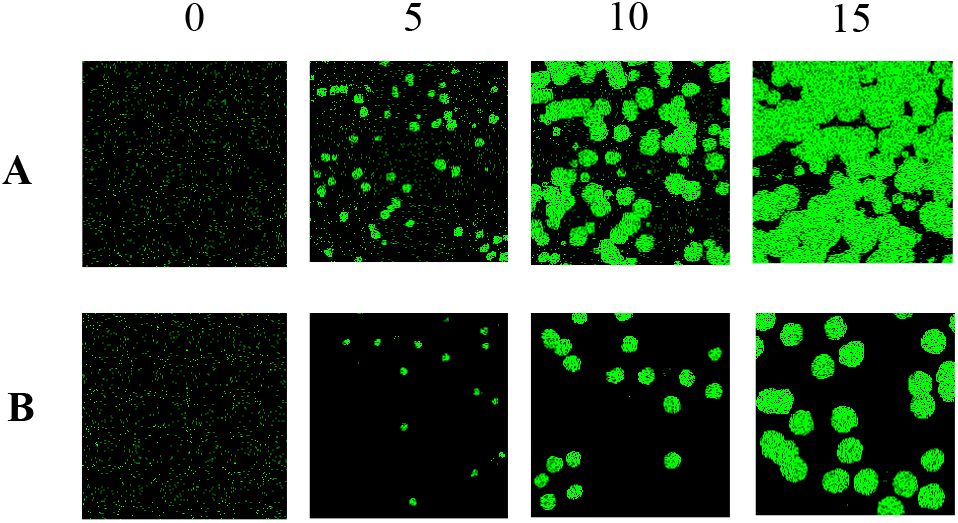
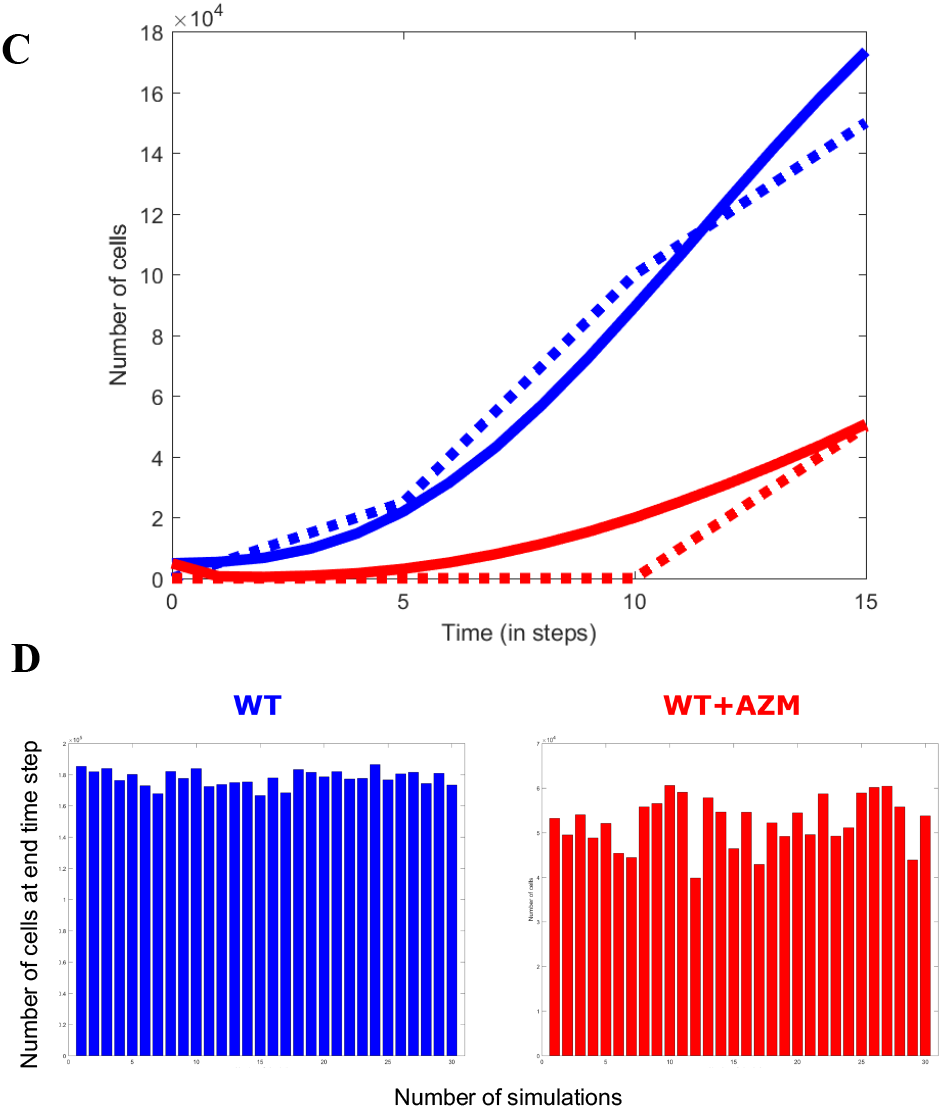
Simulations of spatiotemporal evolution of *P. aeruginosa* using CA model. 2-D top view for 15 time steps, (A) WT and (B) WT+AZM. (C) Cell growth in time WT (blue) and WT+AZM (red), dotted lines indicate experimental profiles. (D) Cell growth number at 15th time step for 30 independent simulations, WT (left, blue) and WT+AZM (right, red).

#### 3.1.2 Cellular Automata Model Rules

Our previous cancer CA model had rules that were modified from Conway’s game of life. The rules, although abstract or oversimplification, can generate complex self-organizing spatiotemporal patterns that have been explored in numerous scientific fields. Our intention here is to first use these popular simple rules and gradually modify them to find a suitable set of rules for *P. aeruginosa*.

We adopted a similar approach where we began the model with rules:

i. Any immotile cell with less than *X_1_* live neighbors dies, caused by under-population
ii. Any immotile cell with *X_2_* or *X_3_* live neighbors becomes motile cell on to the next generation
iii. Any immotile cell with more than *X_4_* neighbors dies, caused by overcrowding
iv. Any dead/empty cell with *X_5_* to *X_6_* live neighbors becomes live cell as by reproduction (division)
v. Any motile cell moves randomly to another empty space in time
vi. Any motile cell that cannot move becomes immotile cell on the next generation

where for Conway’s game of life, *X_1_* = 2, *X_2_* = 2, *X_3_* = 3, *X_4_* = 3, *X_5_* = 3. Here we will fit *X_1_* to *X_5_* using genetic algorithm with the experimental growth numbers in Figure 1. Note that we have introduced additional rules 5 and 6 to consider movement of single cells in time, since planktonic single *P. aeruginosa* contain polar flagellum which allows them to be motile.

## 4. Results

Figure 2A-C shows the simulations of our CA model fitted to *P. aeruginosa* growth in Figure 1. Basically, we were required to fit *X_1_* to *X_6_* separately for the WT and WT+AZM condition. We performed hundreds of simulations, using the aid of genetic algorithm to fit the data [13]. Notably, the main difference between the 2 models pointed to only one key ‘parameter’: the percentage of moving cells (Table 1). In other words, according to our simulations, cell movement is repressed by the antibiotic AZM resulting in slower growth rates.

**Table 1.**
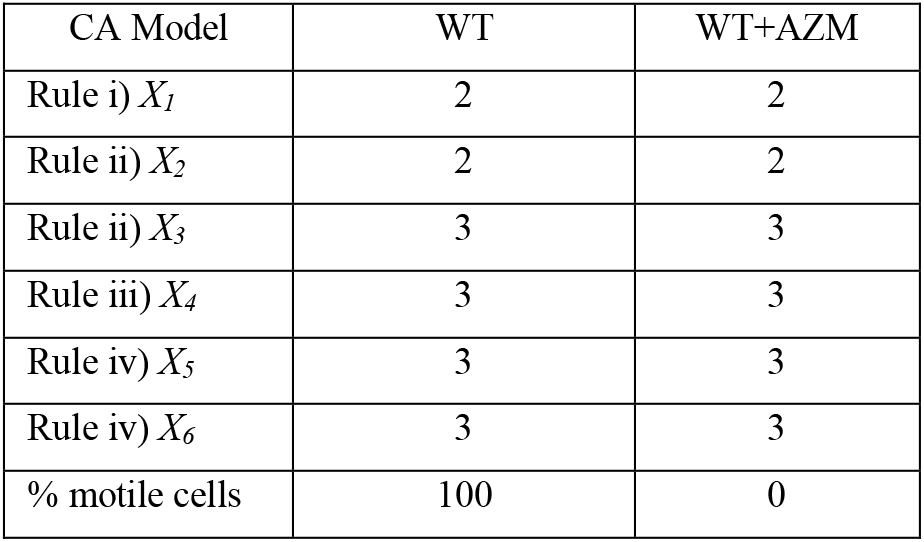
CA model parameters for fitting WT and WT+AZM

We also simulated the final outcome for 30 simulations to check the effect of variability due to random orientation of initial cell distributions. Figure 2D shows the random positioning on average supports the overall experiments.

Using the WT and WT+AZM fitted models, we subsequently simulated the longer term effect on the cell numbers. Notably, after 60 time steps, both model converges to the same cell numbers (Fig. 3). This result is reminiscent of the growth curves shown by another work by Häussler and colleagues [14], who used the same protocol as Gillis and Iglewski [6].

**Fig. 3.**
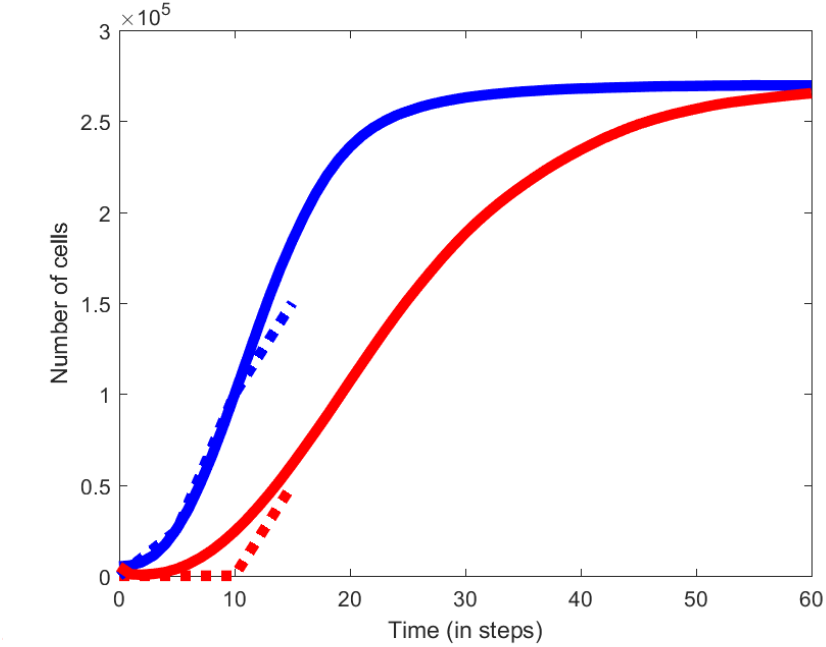
Longer term CA model simulations of spatiotemporal evolution of *P. aeruginosa* using WT (blue) and WT+AZM (red).

Next, we investigated, from the rules, which one is key for suppressing the resurgence of cell proliferation and biofilm formation. After several considerations, we found that *rule* 4 parameters are crucial for suppressing cell proliferation (Fig. 4). In other words, our model proposes the development of drugs that would be able to regulate the dispersion of cells or that prevents cell cluttering. That is, according to *rule* 4, more empty cells between cells prevent cell to cell contact, thereby, is crucial for controlling biofilm formation.

**Fig. 4.**
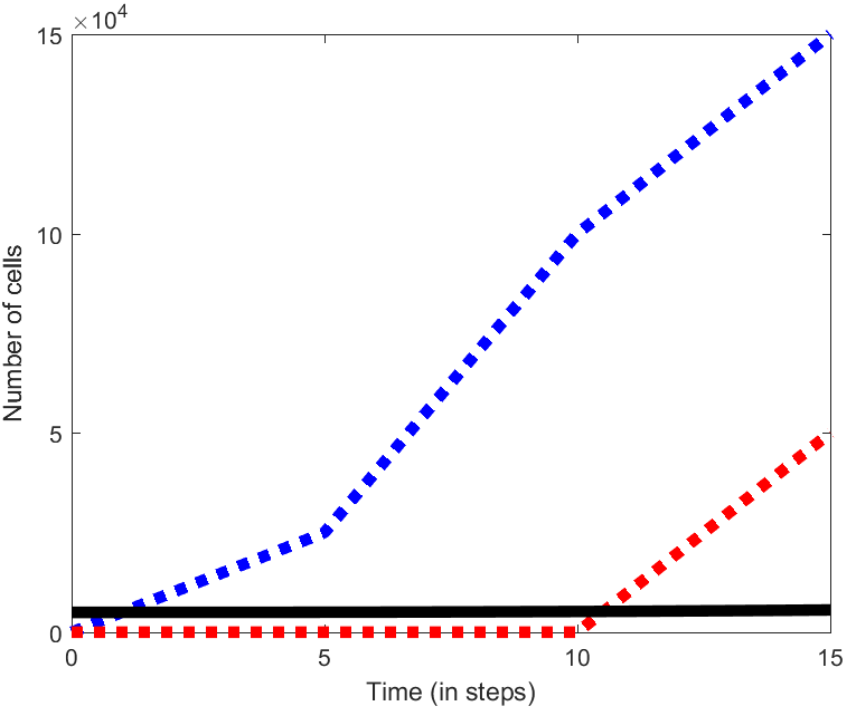
Simulations of spatiotemporal evolution of *P. aeruginosa* using a modified (*rule* 4, *X_5_* = 5 and *X_6_* = 6) CA model that prevents biofilm formation.

## 5. Discussion

In this paper, we have developed a discrete CA model to understand the spatiotemporal self-organizing patterns of *P. aeruginosa* biofilm. The initial rules were taken directly from the famous Conway’s game of life with an additional rule included to factor single cell flagellar random movement. The parameters of the model were fitted with experimental profiles available for the biofilm growth for two conditions. As a result, we developed a CA model, with only one parameter separating WT and WT+AZM simulations.

Notably, our model simulations not only recapitulate the growth profiles of both the untreated and treated biofilm successfully (Fig. 1C and 2C), they also capture the spatial organization of the cells/biofilm over time (Fig. 1A-B and 2A-B). The model predictions suggest that adding the antibiotic agent inhibits the movement of certain single planktonic *P. aeruginosa* which retards their growth. However, subsequently, the inhibition succumbs due to the other rules (1 and 4) to form delayed biofilm. Thus, our model predictions indicate that AZM, on top of killing the cells, is also regulating the cell movement mechanisms such as those involved in flagellar functioning. This delays the biofilm progression.

Experimentally, although AZM is shown to suppress *P. aeruginosa* biofilm [6,14], its mechanism of action still remains poorly understood. We next searched the literature on high-throughput transcriptomics and proteomics related works on AZM treated *P. aeruginosa*. Remarkably, we found the work by Häussler and colleagues [14] supporting our model prediction. In their work, they have shown that the genes and proteins related to flagellar are indeed down-regulated using the same dosage of AZM treatment compared with WT.

Moving further, to find a condition that would effectively suppress *P. aeruginosa* biofilm formation, we searched for the most appropriate rules and their parameter values. The best model suggests that *rule* 4 should have parameter values *X_5_* = 5 and *X_6_* = 6, which will prevent biofilm formation and keep the cell numbers almost unchanged throughout time (Fig. 4). It will be interesting and crucial to identify the biological target that will regulate *rule* 4 by preventing cell to cell contact. Trying adhesins inhibitors with AZM may be a viable next option.

In summary, our work here highlights the need for interdisciplinary research to understand and combat the complexities of living systems, such as controlling the pathogenic microorganisms that endanger the lives of infected people. However, further work is required to validate our final prediction or to test multi-species quorum-sensing bacteria evolution, which is usually a major concern in chronic infection.

## Acknowledgments

The authors thank BioTrans for funding (IAF-PP) the work, N. Lindley and Y. Kanagasundram for discussion.

